# Diet-induced plasticity of life-history traits and gene expression in outbred *Drosophila melanogaster* population

**DOI:** 10.1101/2023.09.02.556027

**Authors:** Akhila Mudunuri, M Chandrakanth, Soumen Khan, Chand Sura, Nishant Kumar, Sudipta Tung

**Author notes:** Equivalent contribution (M Chandrakanth, Soumen Khan). Centre for the Advanced Study of Collective Behavior, University of Konstanz, Konstanz, Germany 78464 International Max Planck Research School for Quantitative Behavior, Ecology and Evolution, Konstanz, Germany 78464 Department of Biology, University of Konstanz, Konstanz, Germany 78464 (Akhila Mudunuri). Epigenetics Department, The Babraham Institute, Cambridge, United Kingdom, CB22 3AT (Soumen Khan). [ORCID 0000-0002-3201-3303] (Sudipta Tung). **Name and address of the corresponding author:** Sudipta Tung DBT/Wellcome Trust India Alliance Early Career Fellow Department of Biology, Ashoka University Plot No. 2, Rajiv Gandhi Education City, Post Office Rai, Near Rai Police Station, Sonipat, Rai, Haryana, India 131029. **Conflict of Interest statement** The authors declare no conflict of interest. **Author contributions:** AM, CM and ST conceived the ideas and designed methodology of the life-history assays in this study; AM, CM, CS, NK, ST collected the data; AM analysed the data. SK and ST conceived the ideas and designed methodology of the transcriptomics part in this study; SK collected the data; SK analysed the data. AM, SK and ST led the writing of the manuscript. All authors contributed critically to the drafts and gave final approval for publication. **Data availability statement** Life-history data will be available on Dryad Digital Repository upon acceptance of the manuscript. The raw data for the RNAseq experiment is submitted in GEO repository, and can be accessed using the link https://www.ncbi.nlm.nih.gov/geo/query/acc.cgi?acc=GSE241397.

## Abstract

Food is fundamental for the survival of organisms, governing growth, maintenance, and reproduction through the provision of essential macronutrients. However, access to food with optimum macronutrient composition, which will maximize the evolutionary fitness of an organism, is not always guaranteed. This leads to dietary mismatches with potential impacts on organismal performance. To understand the consequences of such dietary mismatches, we examined the effects of isocaloric diets varying in macronutrient composition on eight key organismal traits spanning across the lifespan of a large outbred *Drosophila melanogaster* population (n ∼ 2500). Our findings reveal that carbohydrate-reduced isocaloric diets correlates to accelerated pre-adult development and boosts reproductive output without impacting pre-adult viability and body size. Conversely, an elevated dietary carbohydrate content correlated to reduced lifespan in flies, evidenced by accelerated functional senescence including compromised locomotor activity and deteriorating gut integrity. Furthermore, transcriptomic analysis indicated a substantial difference in gene regulatory landscapes between flies subject to high carbohydrate vs high protein diet, with elevated protein levels indicating transcriptomes primed for reduced synthesis of fatty acids. Taken together, our study helps advance our understanding of the effect of macronutrient composition on life history traits and their interrelations, offering critical insights into potential adaptive strategies that organisms might adopt against the continual dietary imbalances prevalent in the rapidly evolving environment.

## 1. Introduction

The fundamental necessity of an organism is food. It provides all the essential resources and nutrients essential to the survival, growth, maintenance and reproduction. Central to these resources are macronutrients, utilized in distinct yet interconnected functions in an organism’s physiology. For instance, proteins primarily serve as building blocks for growth and repair, while carbohydrates and lipids are generally utilized to fuel the energy requirements for cellular and organismal functions (Jequier, 1994; Wu et al., 2014). Therefore, a balanced diet comprising of optimal macronutrient composition is key to maximize the evolutionary fitness of an organism (Altmann, 1991; Brunner et al., 2001; Hawley et al., 2016).

However, the availability of such an optimal diet is not always guaranteed in natural settings, leading potentially to dietary mismatches (Gurven & Lieberman, 2020; Logan & Jacka, 2014; Wiggins & Wilder, 2018). The consequences of this dietary shifts on an organism’s performance and life history traits are difficult to predict based on our current knowledge. Furthermore, a comprehensive study of diet’s impact on an organism’s performance is difficult, given that it requires exact manipulation of food components and an understanding of their short-term as well as long-term effects on performance. Therefore, conducting such studies in natural settings and using organisms with longer lifespans can be challenging. Additionally, considering that diet may simultaneously influence multiple organismal traits, an effective study requires a system where these traits are well-characterized. It is also crucial to understand how these nutrients are being metabolised, which lead to the changes in the observed phenotypes. In this context, *Drosophila melanogaster* serves as a suitable model system for investigating diet’s influence on organismal traits spanning across its life stages. Given its well-understood laboratory ecology, well-characterized traits, and a comparatively short lifespan, *Drosophila* allows for precise dietary component manipulations and longitudinal performance tracking in controlled laboratory settings (Carvalho et al., 2012; Catterson et al., 2010; Jang & Lee, 2018; Piper et al., 2005; Skorupa et al., 2008).

Historically, in studies correlating diet’s influence to organismal performance, the calorific value of diet has gathered considerable attention given the impact of dietary restriction on lifespan across various taxa, including invertebrates (Houthoofd et al., 2005; Jiang et al., 2000; Partridge et al., 2005), rodents (McCay et al., 1935), and humans (Holloszy & Fontana, 2007). However, recent findings highlight that diet’s impact is more nuanced, with isocaloric diets varying in macronutrient composition significantly affecting organismal life history including lifespan (Mair et al., 2005; Shi et al., 2021). While all macronutrients are essential, empirical evidence highlights the critical importance of dietary composition, often depicted as the ratios of macronutrients, for organismal growth, reproduction, and survival (Raubenheimer & Simpson, 1997; Simpson & Raubenheimer, 2009).

However, in-spite of several efforts, knowledge about the precise role of macronutrient composition in diets in influencing organismal performance is still discrepant (Bruce et al., 2013; Lee, 2015; Lee et al., 2008; Lushchak et al., 2012, 2014). These discrepancies across the existing published reports can be attributed to several interconnected reasons. First, a standardized method of varying nutrient composition across reports does not exist, with studies being varied in the choice of ingredients and thereby often introducing confounding effects due to changes in caloric content. Second, many studies examine the impact of diet on a specific trait or a limited set of traits, potentially leading to inconsistencies in the interpretation of trait correlations. Third, the reports often vary in terms of the timing of dietary change implementation relative to the organism’s life stage. And finally, strong interaction of diet with genetics (G×E interaction) (Heianza & Qi, 2017; Roche et al., 2005; Ussar et al., 2015) complicates the synthesis of results from different experiments, particularly when most existing studies use inbred *Drosophila* lines each with diverse genetic backgrounds. Taken together, these shortcomings highlight the need for a consistent experimental framework to comprehend the effects of diet on traits across various life stages.

To circumvent such challenges, in this study we examined the effect of diet-composition-induced plasticity across eight key life history traits spanning various life stages, thereby aiming to obtain a holistic understanding of the impact of diet on organismal fitness. Using a well-characterized outbred population of *Drosophila melanogaster* (Sarangi et al., 2016) and sampling a large number of individuals during assays, we ensured that our results are free from biases of a specific genotype or a particular inbreeding trajectory. We have utilized isocaloric diet regimes with five different protein-to-carbohydrate ratios to control the confounding effect of diet’s calorific value. Our findings suggest a positive correlation between increased carbohydrate content in diet to delayed pre-adult development, compromised reproductive output, and a shortened lifespan. In addition, using transcriptomic studies, we observe significant differences between gene expression profiles between flies fed on different dietary conditions, with aberrant transcriptional upregulation of genes involved in muscle development, being observed in flies fed on diet with higher carbohydrate content. Our results provide crucial insights into how macronutrient composition shapes various life history traits and their correlations and help advance our understanding of how organisms might adapt to dietary fluctuations in today’s rapidly changing world.

## 2. Materials and Methods

### 2.1 Maintenance regime of fly populations

The flies used in this experiment were derived from a large (n = ∼2100), outbred Baseline Population (BP). This baseline population was derived by equally mixing MB_1-4_ (Melanogaster Baseline) populations, whose detailed ancestry can be found in a previous articles (Sarangi et al., 2016; Shenoi et al., 2016). The BP population was maintained at 25°C in constant light, following a 14-day generation cycle for over 30 generations before the current experiments. Throughout this period, the flies were given a baseline diet composed of corn, sugar, and yeast (recipe provided in Table S1). This diet had a calorific value of 714kcal/L and a protein-to-carbohydrate ratio (henceforth, P:C) of 0.4. For routine population maintenance, we collected ∼300 eggs in each of seven clear Laxbro^©^ FLBT-20 plastic bottles, each containing ∼50ml of food. On the 11^th^ day post egg-collection, the adults were transferred to a large plexi-glass cage (25 cm × 20 cm × 15 cm). After two days, a fresh petri-plate containing baseline media was provided for about 14-hour egg-laying period. On the 14^th^ day post egg-collection, these eggs were then transferred into seven bottles following the above method to initiate the next generation. The adult flies remained in the cage as a backup until egg collection for the subsequent generation.

### 2.2 Diets

To evaluate organismal performance across different macronutrient compositions, we formulated five isocaloric diets (matching the calorific value of the baseline diet) with distinct P:C ratios of 0.2, 0.25, 0.4, 0.55, and 0.7. These ratios were adjusted by modifying the amounts of sugar and yeast in the food, while maintaining consistent caloric values (see Table S1 for exact recipes). The diet with a P:C ratio of 0.4 matches the baseline, P:C ratio of 0.2 represents the maximum carbohydrate and minimum protein, and P:C ratio of 0.7 is the converse. It is worth noting that free amino acid or protein powder is often preferred over yeast for dietary protein manipulation in flies (Chang et al., 2001; Fanson & Taylor, 2012; Maklakov et al., 2008; Piper et al., 2014; although see Fanson et al., 2009; Lee et al., 2008) due to complex composition of yeast beyond just protein. In our study, we have also accounted for the carbohydrate content of yeast to accurately calculate the P:C ratio of experimental diets (see Table S1).

### 2.3 Assays

For all assays, 100-150 eggs from the baseline population were collected in plastic bottles containing 50 ml of the respective food types to generate the experimental flies. Throughout the study, we ensured consistent environmental conditions, maintaining constant light, a temperature of 25°C, 80–90% humidity, abundant nutrition, and an uncrowded rearing environment. Chronological age of the flies was also uniformly controlled across assays. Our investigation focused on measuring four sets of important life-history traits related to the flies’ preadult development, reproductive output, lifespan, and functional senescence across the aforementioned five dietary regimes.

#### 2.3.1 Preadultdevelopment and survivorship assay

For this assay, exactly 100 stage-synchronized (Prasad et al., 2000) eggs were collected from the baseline flies in each of five replicate bottles containing 50 ml food of a specific experimental regime. The same is repeated for all the five experimental food regimes. The bottles were maintained under the environmental conditions, which were identical to the baseline population. After the pupae had darkened, the bottles were regularly checked at 2-h intervals and the number of eclosing males and females was recorded against the corresponding eclosion time. We continued these observations until no flies eclosed for over 48h from a bottle. From this data, we computed average time to eclosion, preadult survivorship (i.e. the percentage of flies eclosed out of 100 stage-synchronized eggs) and sex ratio (i.e. the number of males divided by the number of female flies eclosed) of the emerged adults for each of the five replicate bottles in each treatment regime and subjected to statistical analysis.

#### 2.3.2 Fertility assay

To understand the effect of diet on reproductive output, we examined the number of viable offspring produced by the females reared on different diet regimes. We generated experimental flies by collecting eggs and subsequently raising the eclosed adults in mixed-sex groups under five distinct experimental treatment diets. On the 12^th^ day post-egg collection, we introduced two pairs of male and female flies into each vial containing 6ml of the respective treatment diet. These flies were given 24 hours to lay eggs, after which the adults were removed. This process was replicated in 30 vials for each dietary treatment. We then calculated per capita female fertility based on the number of adult offspring that emerged from each vial. Since the diet treatment has no effect on fly viability (see Results and Figure S3), the data obtained would be primarily due to the effect of diet on fertility. To determine whether variations in fertility data correspond to differences in body size across the dietary regimes, we used wing length of adult flies as a proxy for body size (see Supplementary Text S1 for details).

#### 2.3.3 Lifespan and functional senescence assays

Subsequently, we investigated the effect of nutrition on the fly physiology as the fly ages. To check this, we measured lifespan, and quantified male activity and female gut health at old age to test the state of these functional traits in flies reared under different diet regimes. For measuring lifespan, on the 12^th^ day from the day of egg collection, we separated the experimental flies by sex under light CO_2_ anesthesia and placed them in same-sex groups of 8 individuals into glass vials containing 6mL food of respective diet regime. We conducted 10 such replicates per sex for each of the five diet regimes. We monitored the vials daily at a particular time (arbitrarily set at 1200 noon) and they were transferred into fresh food vials every alternate day, till the last individual died. The number of days one fly lived from the time of the assay setup is considered its lifespan. Average lifespan was computed for each replicate setup and was subjected to statistical analysis.

To quantify the senescence status of activity, we monitored locomotor activity of 34-day old male flies using Drosophila Activity Monitoring (DAM) Systems. We recorded their activity for 1 hour 10 minutes. For analysis, the first 10 minutes of data was excluded and the next 1 hour of data was used to calculate fly activity. Activity of 32 male flies were measured per diet regime. As a control to ensure that the effects of aging are due to the rearing diet, we measured locomotor activity of 12-day old male and female flies using DAM Systems with the same protocol as above. Activity of 32 males and 32 females was collected and subjected to statistical analysis.

To check the gut health, 34-day old female flies were put in a cohort of 10 individuals into glass vials with approximately 2 ml of fly food colored with 0.07 ml of brilliant blue dye (Arun Colour Chem Pvt. Ltd.) at a concentration of 2.5% (wt/vol). 10 such replicate vials were made per diet regime. The flies were allowed to stay in the vial for 1 day and then they were checked under the microscope for a Smurf phenotype (Martins *et al*. 2018). The number of flies which turned Smurf was recorded to compute the proportion of Non-Smurf flies for each vial. We computed arcsine of the square root of the proportions (following Zar, 1999) and subjected to statistical analysis.

#### 2.3.4 RNA extraction and Library Preparation

On the 13^th^ day post egg collection, 12 female flies were collected for a particular biological replicate and snap frozen in Trizol solution. Three such replicates were made for each of the diet regimes with P:C ratios of 0.25, 0.4, 0.55 and 0.7. RNA isolation was performed using standard Phenol Chloroform extraction methods and RNA quality determined on Agilent Bioanalyzer System 4000. RNA libraries were performed post polyA enrichment using the NEBNext Ultra II RNA Library Prep Kit for Illumina, using standard protocol. Libraries were sequenced using a 150BP paired end chemistry on Novaseq6000.

### 2.4 Statistical and bioinformatic analysis

#### 2.4.1 Analysis of life-history traits

First, we evaluated how different diet regimes affected the average developmental time of the flies. The data for average developmental time, which is continuous and unbounded (see Section 2.3.1 for details), was analyzed using a two-way full-factorial ANOVA. In this analysis, both diet—specified by protein-to-carbohydrate (P:C) ratios of 0.2, 0.25, 0.4, 0.55, and 0.7—and sex (Male or Female) were treated as fixed factors. Preadult survivorship data being percentage values, it was arcsine square root transformed prior to analysis. In order to test the extent of diet-induced plasticity, transformed preadult survivorship data and sex ratio data were subjected to one-way fixed factor ANOVA with diet (P:C ratios of 0.2,0.25,0.4,0.55,0.7) as the fixed factor. A post-hoc analysis was done using Tukey’s HSD test. All the above statistical analysis was done using STATISTICA^®^ v5 (StatSoft Inc., Tulsa, Oklahoma).

Subsequently, we examined the effect of dietary conditions on fertility per female, treating it as count data. For the analysis, we employed a Negative Binomial Generalized Linear Model (GLM) with a log link function, using the ‘glm.nb’ function from ‘MASS’ package in R version 4.3.1. The ANOVA table was generated with the ‘Anova’ function from the ‘car’ package. To identify specific differences between the levels of the diet regimes (*i.e.* P:C ratios of 0.2, 0.25, 0.4, 0.55, 0.7), we conducted post-hoc analysis using Tukey’s HSD test, implemented via the ‘glht’ function in the ‘multcomp’ package.

We analyzed the effects of Food, Sex, and their interaction on wing length, measured as a proxy for body size, using a GLM with a Gaussian family and identity link function. The ANOVA table was generated using Type III sums of squares. Post-hoc comparisons among Food levels were conducted using Tukey’s HSD test. All analyses were performed in R version 4.3.1 with the ‘glm’ function for model fitting, the ‘Anova’ function from the ‘car’ package for the ANOVA table, and the ‘glht’ function from the ‘multcomp’ package for Tukey’s test.

To test the impact of diet regimes on lifespan, we subjected the average lifespan data to a two-way full-factorial ANOVA with diet (P:C ratios of 0.2, 0.25, 0.4, 0.55, and 0.7) and sex (Male or Female) were treated as fixed factors. A post-hoc analysis was done using Tukey’s HSD test. This statistical analysis was done using STATISTICA^®^ v5 (StatSoft Inc., Tulsa, Oklahoma). Further we compared the survivorship curves using a Cox Proportional-Hazards model. The ‘Surv’ function from the ‘survival’ package in R version 4.3.1 was used to create the survival object, incorporating both time and event status. Food levels (P:C ratios of 0.2, 0.25, 0.4, 0.55, and 0.7) were set as factors and P:C ratio of 0.4 was considered as the reference category. The model was fitted using the ‘coxph’ function and summary statistics were obtained. Hazard ratios for each factor and interaction term were visualized using the ‘ggforest’ function from the ‘survminer’ package, and survival curves for both sexes across different food levels were plotted using ‘ggsurvplot’ from the ‘survminer’ package and customized with ‘ggplot2’.

We used a negative binomial GLM to analyze the effects of Food and Sex on first-hour locomotor activity levels of the young flies using a full factorial model in R version 4.3.1 using the ‘glm.nb’ and ‘Anova’ functions from the ‘MASS’ and ‘car’ packages, respectively. For analyzing activity data of old male flies, we used a zero-inflated negative binomial model, as it provided a better fit according to Vuong’s closeness test. The model was implemented in R using the ‘zeroinfl’ function from the ‘pscl’ package. Subsequently, we calculated Cohen’s d effect sizes to quantify the differences in activity levels across diets. Effect sizes were categorized as small (0.2 < d < 0.5), medium (0.5 < d < 0.8), or large (d > 0.8) following Cohen’s guidelines (Cohen, 1988).

The effect of diet on gut integrity in aging female flies was evaluated using a GLM with a Gaussian family and an identity link function. The percentage of females exhibiting the ‘Smurf’ phenotype was arcsine square root-transformed prior to analysis. Post-hoc pairwise comparisons were conducted using Tukey’s HSD test. All analyses were performed in R version 4.3.1. The graphs were made using ggplot2 (Wickham, 2016) package in R version 4.3.1 and RStudio v2022.07.1 (RStudio Team, 2020).

#### 2.4.2 RNA-seq analysis

Raw FASTQ files were subject to initial quality control analyses using the FASTQC software v0.12.1 (Andrews, 2010). FASTQ files were then trimmed using Trim-galore v0.6.10 (Krueger, 2015) and aligned to the *Drosophila* genome (dm6) using hisat2 (Kim et al., 2019). The Seqmonk software v1.48.1 (https://www.bioinformatics.babraham.ac.uk/projects/seqmonk/) was used to generate raw counts and differential gene expression performed using DESeq2 (Love et al., 2014). PCA and volcano plots were generated in R using ggplot2. Genes were first filtered on padj<0.05 and Fold-change (FC)>1.5 and sorted based on FC. The top 40 differentially expressed (DE) genes between diets with a P:C ratio of 0.4 and P:C ratio of 0.25, P:C ratio of 0.4 and P:C ratio of 0.7 and the top 5 DE genes between P:C ratio of 0.4 and P:C ratio of 0.55 were selected, and unique genes amongst the list of 85 genes (final total of 77 genes) were used for heatmap analyses. The g:Profiler software (Raudvere et al., 2019) was used for Gene Ontology analysis using a custom background (all expressed genes) and Bonferroni padj<0.05 was used as the cutoff for significant Gene Ontology-terms.

**Table1.** Replication Statement.

| Experiment | Scale of inference | Scale at which the factor of interest is applied | Number of replicates at the appropriate scale |
| --- | --- | --- | --- |
| Preadult developmental time assay | Population | Bottle (with 100 eggs in each) | 5, 5, 5, 5, 5 |
| Pre-adult viability assay | Population | Bottle (with 100 eggs in each) | 5, 5, 5, 5, 5 |
| Sex ratio assay | Population | Bottle (with 100 eggs in each) | 5, 5, 5, 5, 5 |
| Body size assay | Individual | Individual sex | 30, 30, 30, 30, 30 |
| Fertility assay | Individual | Individual pair (one male and one female) | 60, 60, 60, 60, 60 |
| Lifespan assay | Male and Female population | Vial (with 10 individuals of one sex in each) | 10, 10, 10, 10, 10 (vials per sex) |
| Locomotor activity assay of young flies | Male and Female population | Individual | 32, 32, 32, 32, 32 (individuals per sex) |
| Locomotor activity assay of old male flies | Individual | Individual | 32, 32, 32, 32, 32 |
| Smurf assay with old female flies | Population | Vial (with 10 individuals in each) | 10, 10, 10, 10, 10 |
| RNAseq | Population | Group of 12 individuals | 3, 3, 3, 3 |

## 3. Results

### Reduced carbohydrate proportion in isocaloric diet accelerates developmental speed without affecting pre-adult survivorship or sex ratio

To understand how dietary composition affect organismal performance, we first looked into developmental timing, *i.e.* the time taken for an egg to develop into an adult *Drosophila*, when subject to different diets. We report a significant effect of macronutrient composition of diet on the developmental time in flies (Figure 1, F_4,38_ = 79.67, *p* < 0.0001). Our results indicate that flies reared on an isocaloric diet with a P:C ratio of 0.7 were found to develop faster compared to flies reared on a diet with a P:C ratio of 0.2 (Tukey’s *p* = 0.0001), 0.25 (Tukey’s *p* = 0.0001), and marginally faster than the flies reared on P:C ratio of 0.4 (Tukey’s *p* =0.057). Remarkably, despite a minor decrease in dietary carbohydrate content, flies reared on a diet with a P:C ratio of 0.4 exhibited significantly accelerated development compared to those on a P:C ratio of 0.25 (Tukey’s *p* = 0.0001). Similar to previous reports, we too observe that female flies eclose early in comparison to males (F_1,38_ = 7.79, *p* = 0.008), however, the food x sex interaction was not significant (F_4,38_ = 0.57, *p* = 0.687). The same pattern is also visible when the data is visualized through the percentage-eclosion-across-time curves (Figure S1). Overall, our findings suggest that, despite equal calorific values, the relative proportions of dietary carbohydrates and proteins play a critical role in the egg to adult developmental time of *Drosophila melanogaster*. Specifically, we report that reducing carbohydrate content and increasing protein content in the diet is positively correlated to pre-adult developmental rate. We further found that pre-adult survivorship (Figure S2, F_4,19_ = 0.528, *p* = 0.716) and the sex ratio of the eclosing adults (Figure S3, F_4,19_ = 0.958, *p* = 0.452) were comparable across the experimental diets.

**Figure 1.**
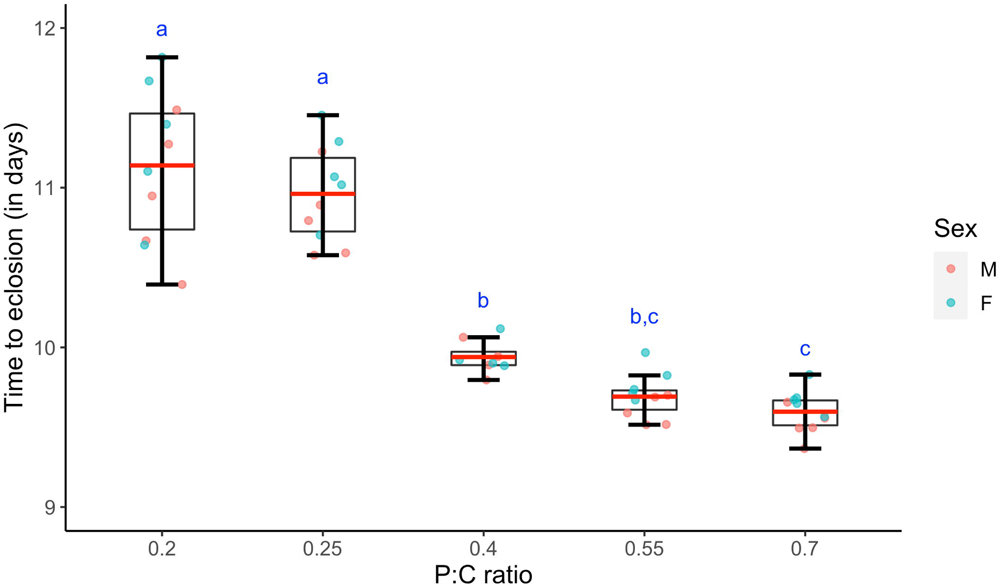
Effect of diet on development. Mean (±SEM) development time of individuals reared on different diets is plotted against the diet which they were reared on. The box plot represents combined average developmental time data for both males and females.. The red line, lower and upper edge of the box, lower and upper end of the whiskers represent the mean, 25th, 75th, 5th, 95th percentile of the development time data. The scatter points denote the mean of the developmental time of the individuals in each bottle. Different lower-case alphabets denote statistically significant differences (*p*<0.05).

### Higher dietary carbohydrate proportions reduce reproductive output without affecting body size

We report significant differences in viable offspring produced per adult pair, composed of one male and one female, in relation to dietary proportion of protein and carbohydrate (Figure 2, F_4,145_ = 61.239, *p* < 2.2e-16). Interestingly, even slight increase in carbohydrate content (and thereby, decrease in protein content) was negatively correlated to the reproductive output, with flies reared on an isocaloric diet with a P:C ratio of 0.2 producing fewer number of viable offspring than flies reared on a diet with a P:C ratio of 0.25 (Tukey’s *p* < 0.001). In contrary, flies reared on increased protein concentrations display higher fertility, with a significant increase seen between diets with protein to carbohydrate ratio 0.25 and 0.4 (Tukey’s p=0.009), and between flies reared on a diet with a P:C ratio of 0.70 and a diet with a P:C ratio of 0.55 (Tukey’s p= 0.04). In terms of wing length as a measure of body size, although the main effect of food was significant (F_4,290_ = 2.678, *p* = 0.03), Tukey’s pair-wise comparison shows flies were comparable across experimental diets (Figure S4), except for those on the lowest protein-to-carbohydrate ratio (P:C ratio of 0.2) were found to be smaller than those reared on P:C ratio of 0.4 (Tukey’s p = 0.024). Taken together, our results reveal a remarkable reproductive plasticity exhibited by *Drosophila* across varying P:C ratios in isocaloric diets.

**Figure 2.**
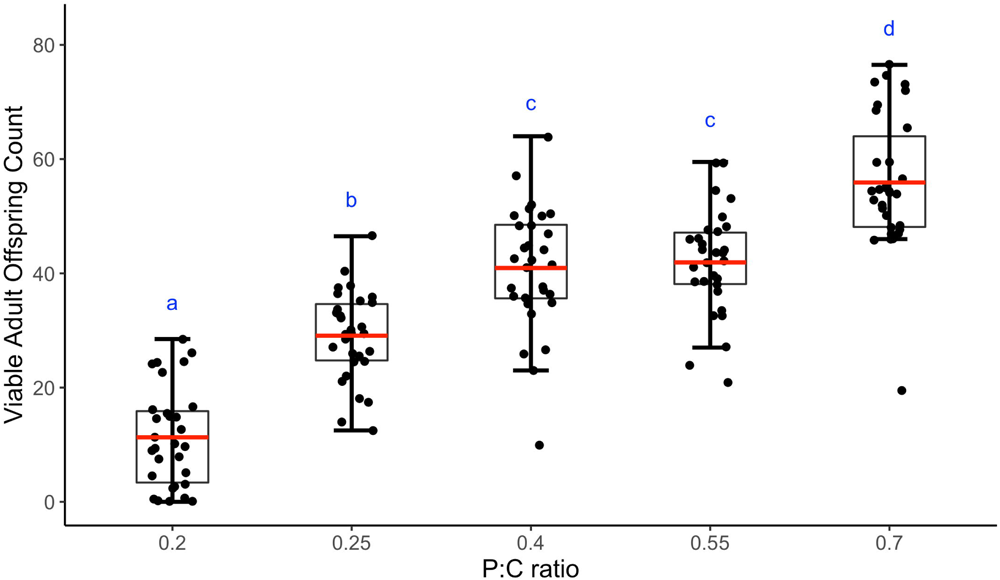
Effect of diet on fecundity. Mean (±SEM) number of viable adult offspring sired by a male-female pair reared on different diets is plotted against the diet which they were reared on. The box plot represents the fecundity of the females reared on diets with different P:C ratios. The red line, lower and upper edge of the box, lower and upper end of the whiskers represent the mean, 25th, 75th, 5th, 95th percentile of the fecundity data. The scatter points denote the number of eggs laid by each female.

### High carbohydrate content in diet shortens fly lifespan

We next evaluated the effect of varying macronutrients on the lifespan of *Drosophila* and observe a significant effect of varying carbohydrate to protein ratio on longevity of flies (Figure 3, F_4,90_ = 35.24, *p* < 0.0001). We observe that a diet with a high P:C ratio of 0.7 prolonged the lifespan of flies significantly when compared to those reared on a diet with a low P:C ratio of 0.2 in both males (Tukey’s *p* = 0.0001) and females (Tukey’s *p* = 0.0001). It is also notable that lifespan of male and female flies was comparable when flies were reared at high P:C ratio of 0.7, whereas with diets having high carbohydrate proportion (P:C ratio of 0.2 and 0.25), female lifespan was substantially lower than that of males (Figure 3). This led to a significant food × sex interaction (F_4,90_ = 11.05, *p* < 0.0001). We observe the same pattern while comparing the entire survival curves (Figure S5) using Cox-proportional hazard model followed by Wald test (Table S9, Figure S6). Overall, our results indicate that given comparable calorific value diets, on one hand we observe a significant and negative correlation of lifespan when the flies are reared on a diet with a lower P:C ratio in diet, and on the other a positive correlation when the flies are reared on a diet with a higher P:C ratio.

**Figure 3.**
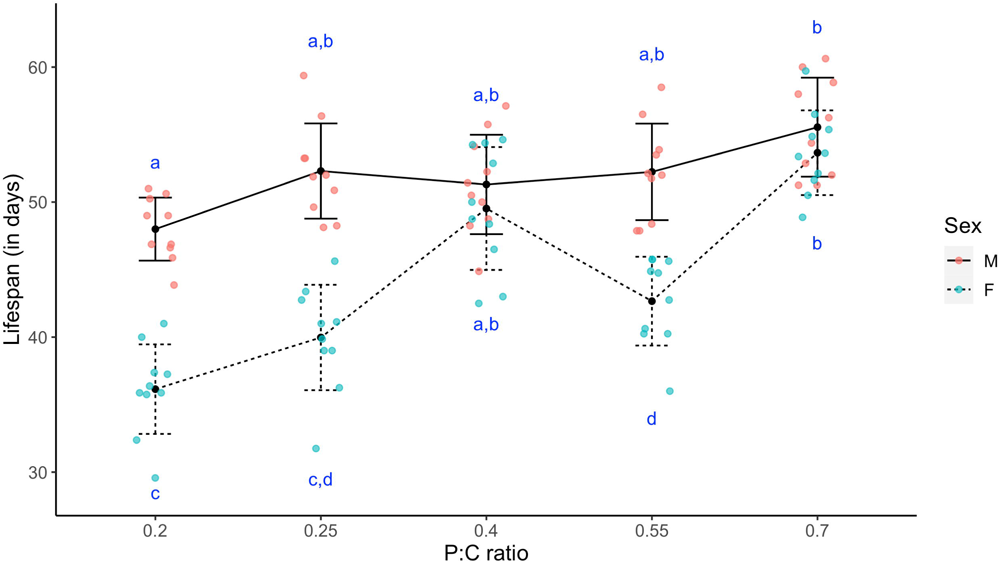
Effect of diet on lifespan. Mean (±SEM) lifespan of the flies reared on different diets. The line-scatter plot represents the average lifespan of flies reared on diets with different P:C ratios The scatter points denote the mean of the lifespan of 10 flies of each sex housed together. Different lower-case alphabets denote statistically significant differences (p<0.05). Different lower-case alphabets denote statistically significant differences (*p*<0.05).

### Carbohydrate accelerates aging and impairs late-life physiology and behavior

Consistent with the effect of macronutrient composition on lifespan, our experiments also reveal the role of varying protein-to-carbohydrate ratios on late-life locomotor activity of males (Figure 4A, F_4,148_ = 2.3621, *p* = 0.05) and gut physiology of female flies (Figure 4B, F_4,45_ = 4.518, *p* = 0.0037). We report that the functional status of both these traits was better in flies that consuming food with higher P:C ratio. Male flies reared on a high protein diet with a P:C ratio of 0.7 were more active than those reared on a high carbohydrate diet with a P:C ratio of 0.25 (Cohen’s *d* = 0.82; Large effect size). Females reared on a low carbohydrate diet with a P:C ratio of 0.7 showed increased gut integrity when compared to females reared on a high carbohydrate diet with P:C ratio of 0.25 (Tukey’s *p* = 0.0043). It’s important to highlight that the observed decline in the performance of the flies under carbohydrate-rich diets are specific to older age. This is because, in their early stages, there was no significant difference in locomotor activity among the flies across any of the diet regimes for either sex (Figure S7; F_4,305_ = 1.7, *p* = 0.15). Our studies, thereby, suggests a correlation between increased dietary carbohydrate to reduced late-life activity in male flies and compromised gut health in female flies.

**Figure 4.**
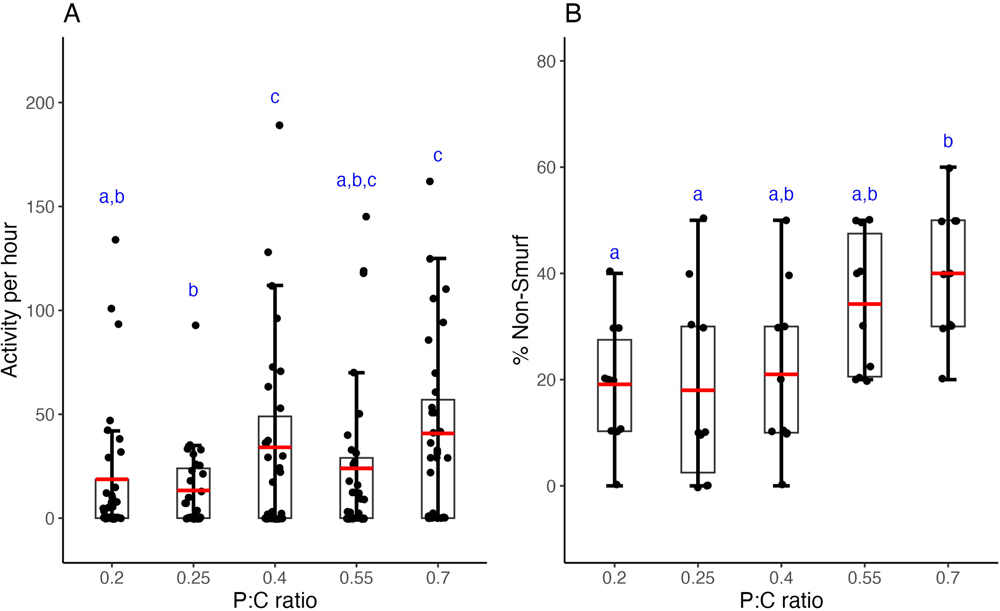
Effect of diet on late-life fitness. (A) Mean (±SEM) locomotor activity of male flies reared on different diets. (B) Mean (±SD) percentage of female flies with intact gut lining. The red line, lower and upper edge of the box, lower and upper end of the whiskers represent the mean, 25th, 75th, 5th, 95th percentile of the data. The scatter points denote the (A) late-life locomotor activity of a male and (B) percentage of females which remained non-smurf (i.e. healthy) in each vial. Different lower-case alphabets denote statistically significant differences (Cohen’s *d* >0.5 or *p*<0.05).

### Impact of protein-to-carbohydrate (P:C) ratios on gene expression levels in *Drosophila melanogaster*

To further investigate the effect of varied protein-to-carbohydrate ratio isocaloric diets on gene expression profiles, we performed transcriptomic analysis of adult female flies. Principal component analysis (PCA) indicates a distinct separation on PC1, between flies fed on diets having a P:C ratio of 0.25 and P:C ratio of 0.7, suggesting highly dissimilar transcriptional profiles (Figure 5A) across the expressed genes (Table S2). Meanwhile, expression profiles of flies fed on diets with P:C ratio of 0.4 and 0.55, while equidistant from the other two conditions (P:C ratio of 0.25 & P:C ratio of 0.7) on the PCA plot, tend to cluster together, thereby indicating that transcriptional profiles of the intermediate diet (P:C ratio of 0.55) mirrors that of the control (P:C ratio of 0.4) (Figure 5A). This is confirmed by the sparse number of differentially expressed (DE) genes (25) between the control (P:C ratio of 0.4) vs P:C ratio of 0.55 diet conditions (Table S4, Figure 5D). On the other hand, we report a total of 160 DE genes in a diet with a P:C ratio of 0.25 and 287 DE genes in a diet with a P:C ratio of 0.7, when compared to the control diet (Figure 5C, Table S3; Figure 5E, Table S5). Interestingly, we find a shift in the regulation profiles between the diet with a P:C ratio of 0.25 and a diet with a P:C ratio of 0.7 when compared to the control, with 95% of the genes being upregulated in the former as compared to 11.8% in the latter (88.2% downregulated) (Figure 5C, 5E). Heatmap analysis of the top 77 DE genes (detailed methodology for choosing the genes provided in the methods section) suggest distinct differences in the gene expression profile across the test groups (Figure 5B). This indicates a rewiring of the gene regulation machinery in response to change in protein-to-carbohydrate ratio of the diet, while also alluding to a dosage-dependent effect for the cause of the phenomenon. To investigate further into the category of genes being differentially expressed in different test groups, we performed Gene Ontology analysis (Table S6, S7). We find that for the diet with a P:C ratio of 0.25 condition (high carbohydrate, low protein diet), genes which are involved in muscle development are upregulated (Figure S8A). Whereas, for the diet with a P:C ratio of 0.7 condition, we find that genes involved in fatty acid synthesis process (including long-chain fatty acyl CoA synthetases), were significantly downregulated, indicating a transcriptome poised for reduced expression of genes involved in synthesis of fatty acids (Figure S8B). Taken together, our study here uncovers a distinct diet-dependent modulation of gene expression profiles in adult female flies.

**Figure 5.**
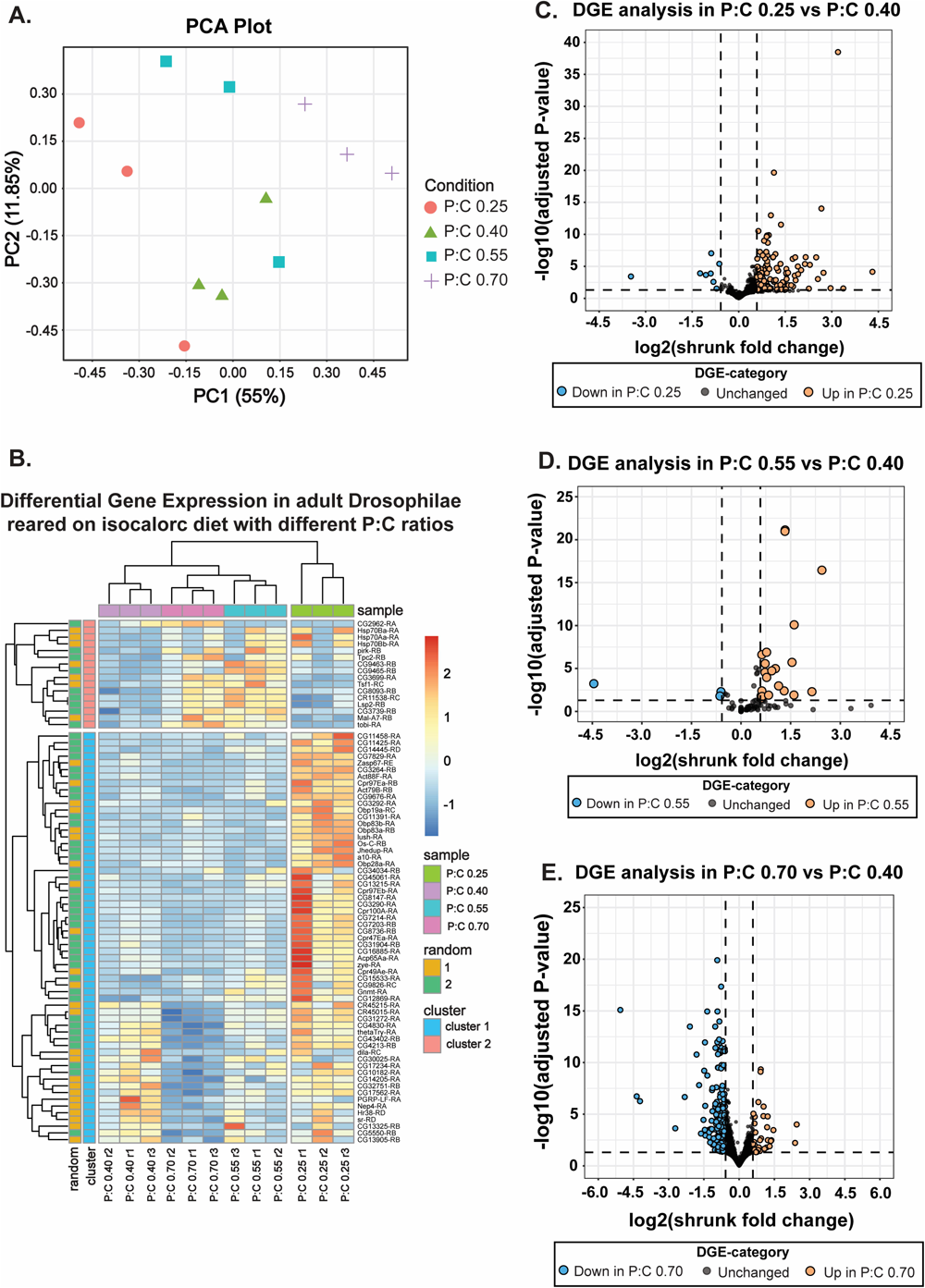
Impact of diet on gene expression profile. (A) Principal Component Analysis (PCA) plot showing the gene expression data of flies maintained on four distinct diet regimes: P:C ratios of 0.25, 0.4, 0.55, and 0.7. (B) Heatmap showing the expression patterns and clustering of the 77 differentially expressed genes from each category. (40 genes between P:C ratio of 0.4 and P:C ratio of 0.25, 40 genes between P:C ratio of 0.4 and P:C ratio of 0.7, and 5 genes between P:C ratio of 0.4 and P:C ratio of 0.55, duplicates excluded). The heatmap color spectrum indicates the normalized z-score values ([z-score= (x-mean(x))/sd(x)], where x represents raw counts). (C, D, E) Volcano plot comparing gene expression levels between flies reared on the baseline diet (P:C ratio of 0.4) and the carbohydrate-rich diet (P:C ratio of 0.25) (C), the intermediate diet with a P:C ratio of 0.55 (D) and the protein-rich diet with a P:C of 0.7 (E). We have applied a 1.5 fold cut-off [log2(shrunk fold change) > abs(0.585)] to identify differentially expressed genes for all three comparisons.

## 4. Discussion

Food provides essential calories for vital activities in an organism, but its significance extends beyond its caloric content (Simpson & Raubenheimer, 2012). Macronutrients obtained from food play a crucial role in growth, development, and maintenance of bodily functions (Raubenheimer & Simpson, 1997; Simpson & Raubenheimer, 2009). However, excessive or inadequate intake of macronutrients can negatively impact organismal health and performance (Lee, 2015; Lee et al., 2008). Our study systematically investigates how altering the relative proportion of carbohydrate and protein in isocaloric diets affects various life-history traits and gene expression profile in *Drosophila melanogaster*. Using a large outbred population (n ∼2100), we explore the complex relationship between diet and organismal performance across multiple phenotypes pertaining to development, reproduction and survival. Previous research primarily used isogenic lines, making it unclear if those results extend to populations with standing genetic variations in natural settings, particularly given substantial genotype × environment interactions observed in other studies (Heianza & Qi, 2017; Roche et al., 2005; Ussar et al., 2015).

In order to examine the effect of diet composition (protein to carbohydrate ratio) on development, we quantified the time taken for flies to develop from embryonic stage to adult eclosion. We observed that an increased proportion of carbohydrates in the diet resulted in a longer developmental period for the flies. This aligns with previous studies that showed a delay in developmental time due to a decrease in protein content (Lee et al., 2008; Matzkin et al., 2011). A possible explanation for this finding can be found in an earlier study by Almeida de Carvalho & Mirth, 2017, who reported that larvae regulate their food consumption in order to achieve an intrinsic threshold level of protein intake. As a result, when raised on an isocaloric high-carbohydrate diet with a low P:C ratio, larvae need to feed for a longer duration to obtain sufficient protein intake, while under a low-carbohydrate diet with a high P:C ratio, larvae reach their protein-intake goal more quickly, resulting in a shorter overall pre-adult developmental time. This hypothesis can be tested by conducting targeted experiments that measure the feeding rate and quantify the amount of carbohydrates and protein consumed relative to the total food intake. Despite a substantial (∼48h) difference in developmental time between flies reared on diets with different compositions, pre-adult viability was not impacted by diet. This suggests that even under the experimental diet with the lowest P:C ratio, flies were able to satisfy their necessary protein requirements for pre-adult growth and development, albeit at a later time point. Additionally, no difference in sex ratio was observed across our experimental diets, indicating that no diet-induced developmental plasticity was present under these dietary regimes, a phenomenon that is often seen in various organisms (reviewed in Rosenfeld & Roberts, 2004).

Pre-adult development time is known to play a crucial role in shaping an organism’s body size and morphology, especially in insects (Atkinson & Sibly, 1997; Davidowitz et al., 2004; Nijhout, 2003; Nylin & Gotthard, 1998). Previous studies have indicated that extending the pre-adult period leads to an increase in body size, while shortening it results in a decrease in body size (Davidowitz et al., 2004). However, in our study, we did not observe a substantial difference in body size, neither in males nor females, across the experimental isocaloric food regimes with varying protein to carbohydrate ratios. This suggests that, as previously explained, the flies reared on a low P:C ratio diet required a significantly longer time to attain their protein requirement. However, in the process, the diet with the lower P:C ratiodid not result in additional growth. It is interesting to note that the flies aim to achieve a certain body size irrespective of the time they take to develop as body size is an important contributor to fitness for the flies in terms of aggression for the males (Partridge & Farquhar, 1983) and higher fecundity for females (Lefranc & Bundgaard, 2000). This illustrates that the correlation between pre-adult development time and body size can be context-specific and dependent on various factors. This raised the question of whether the surplus carbohydrate intake is prioritized for reproductive purposes instead of growth on high-carbohydrate diets (Lee et al., 2008). To test this hypothesis, we investigated reproductive output across the experimental diet regimes.

Interestingly, even though flies reared on different experimental diets had similar body sizes, we observed a significant effect of diet composition on the number of viable offspring produced by adult *Drosophila* pairs (henceforth termed as reproductive output; Figure 3). An increase in the relative proportion of carbohydrates in the diet resulted in a significant decrease in reproductive output. The observed discrepancy of reduced reproductive output on a carbohydrate-rich diet, despite comparable body sizes, aligns with findings from a prior study. This research indicated that elevated sugar intake leads to cholesterol accumulation, subsequently reducing fertility (Brookheart et al., 2017). In addition to the detrimental effect of excessive carbohydrate consumption on reproduction, dietary protein is known to stimulate ovarian stem cell proliferation and oogenesis (Drummond-Barbosa & Spradling, 2001), resulting in greater reproductive output. These findings are supported by several studies that have reported a reduction in female fecundity when dietary protein content is decreased or sugar intake is increased (Lee, 2015; Lushchak et al., 2014; Savola et al., 2021). They also underscore the observed difference in fertility of flies reared on diets with different P:C ratios in the current study, with a staggering more than four-fold increase in the number of offspring observed on the isocaloric experimental diet with the lowest carbohydrate content compared to that with the highest carbohydrate content.

A negative correlation between reproductive output and lifespan is a widely recognized phenomenon across species, including *Drosophila* (Flatt & Kawecki, 2007; Hunt et al., 2004; Rose, 1984). This trade-off is thought to originate owing to resources directed towards reproduction, reducing available resources for somatic maintenance and repair, leading to accelerated senescence and reduced lifespan (Kirkwood & Rose, 1991). However, in a study on *Drosophila*, ablating the germ line had no effect on longevity (Barnes et al., 2006), suggesting that the association between these two traits can be uncoupled (Piper et al., 2017). Interestingly, the association between reproduction and lifespan can even be positively correlated if any factor impacts both the traits in the same manner independently. Carbohydrate-rich diets have been found to reduce lifespan, as well as lower egg-laying capability in fruit flies with low protein diets (Lushchak et al., 2014). Our current study on *Drosophila* also showed that with decreasing P:C ratio, lifespan decreases. Further, it is interesting to note that the signature is more pronounced in females than males, consistent with earlier findings that females are more vulnerable to the adverse effects of high-sugar diets in fruit flies (Chandegra et al., 2017). Supporting the notion of the findings of our study, several studies have demonstrated that a high-sugar diet can lead to cardiomyopathy, elevated oxidative stress, and diabetes-like phenotypes, thereby compromising the lifespan of adult *Drosophila* (Ecker et al., 2017; Musselman et al., 2019; Na et al., 2013; Palanker Musselman et al., 2011), whereas the lifespan of flies increased with increasing protein consumption (Lee, 2015; Savola et al., 2021).

However, our results are in contrast to other studies that found no difference in lifespan between flies reared on high protein and high carbohydrate diets (Watada et al., 2020) or where they found that consuming high carbohydrate and low protein maximizes lifespan (Bruce et al., 2013). There are also empirical evidences suggesting that the effect of dietary P:C ratio on lifespan is non-linear and follows an inverted U-shaped curve (Lee et al., 2008), influenced by factors like genetic backgrounds and temperature. It is understandable that significant changes in parameters, such as increasing the P:C ratio further, altering temperature, or genetic background, could lead to a reduction in lifespan at higher P:C ratios. Given these conflicting findings, a more targeted investigation is needed to understand the mechanistic underpinnings of the impact of macronutrients on lifespan, which is beyond the scope of the current study. Instead, we examined the effects of macronutrient composition on health span by studying the functional senescence of adult locomotor activity and gut physiology.

Lifespan serves as a crucial measure of organismal evolutionary fitness. However, it provides an incomplete binary view of organismal performance. A more informative approach is to examine the gradual decline in functional status with age, or functional senescence, of an organism’s various traits. This perspective is mirrored in human gerontology research, which has recently shifted its focus from extending lifespan to extending the period of healthy, active, and disability-free living, referred to as health span (Partridge, 2016). In our study, we found that male flies reared on a low carbohydrate, high protein diet exhibited greater activity levels at a later age. Similarly in females, with increasing P:C ratio, we observed a delayed decline in gut lining integrity, as indicated by the higher proportion of flies exhibiting the non-Smurf phenotype (Figure 4B). These findings are consistent with prior research, which found that increasing sugar content in the diet increased the proportion of flies displaying a leaky gut phenotype (Pereira et al., 2018, however see Galenza & Foley, 2021). Taken together, our findings suggest that increasing the relative proportion of carbohydrate and decreasing the proportion of protein in the isocaloric diet accelerates functional senescence in flies, leading to a shorter lifespan.

Prompted by the significant influence of diet composition on key life-history traits, including the preadult developmental period, reproductive output, aging, and lifespan, we expanded our exploration to effects at a sub-organismal level. With the understanding that the traits mentioned above are polygenic, we conducted an untargeted transcriptomic analysis of whole-fly samples. The findings reveal a robust diet-induced plasticity in gene expression (Figure 5), corroborating with earlier studies (Haro et al., 2019; Whitaker et al., 2014). Notably, our investigations involved a large, outbred *Drosophila melanogaster* population with substantial standing genetic variation. The consistency across biological replicates attests to the robustness of our data and signal clarity, bolstering the reliability of our results and confidence in the observed gene expression patterns. Gene ontology analysis of differentially expressed genes indicates overexpression of several genes under a high-carbohydrate diet (P:C ratio of 0.25). This overexpression might be attributed to the upregulation of the target of rapamycin (TOR) pathway, a key regulator of multiple transcription factors. TOR pathway is conserved across the species, including *Drosophila* (Tatebe & Shiozaki, 2017), and it is a known downstream target of insulin and insulin-like peptide signaling (IIS) pathway, which is upregulated under high-sugar diet (Teleman, 2009). Intriguingly, an upregulation of genes related to muscle development was also detected (Figure S8A). Consistent with this, past experiments have demonstrated that the TOR signaling pathway can enhance muscle-specific transcription factors in mammalian systems (Risson et al., 2009) and the Insulin receptor/Foxo signaling plays a crucial role in activating muscle protein in *Drosophila* (Demontis & Perrimon, 2009). However, a more detailed investigation of the impact of diet on *Drosophila* muscle stem cells (MuSC) and adult muscle precursors (AMPs) would reveal whether carbohydrate-rich diet is promoting muscle development or the observed plasticity in gene expression is a consequence of elevated maintenance and repair of the muscles in response to damages caused by carbohydrate-rich low-protein diet. Conversely, the gene expression profile of the low-carbohydrate, high-protein diet (P:C ratio of 0.7) reveals a general transcriptome downregulation (Figure 5B, 5C, 5E). Notably, genes implicated in fatty acid synthesis, including long-chain fatty acyl CoA synthetases, which have been linked with obesity and associated metabolic disorders under high fat diet (Bjermo & Risérus, 2010; Sampath & Ntambi, 2011), were also downregulated under high-protein diet (Figure S8B). Although proposed here, a systematic investigation of the precise molecular mechanisms behind these diet-induced transcriptional changes is needed, which is outside the scope of the current study. We recognize that diet impacts may vary across tissues (Ellis et al., 2015), suggesting that future research could focus on a tissue-specific analysis of our findings. Further exploration of the molecular mechanisms triggering these gene expression changes could reveal new pathways and molecular entities involved in dietary adaptations. Ultimately, these findings provide promising avenues for elucidating the molecular basis of dietary adaptations and unraveling the complexities of gene regulation in response to diverse nutrient compositions.

## 5. Conclusion

Our study highlights the significant effects of macronutrient composition on various life history traits - development, reproduction, and survival of *Drosophila*. Specifically, our results indicate that a high proportion of carbohydrates in the diet negatively impacts development, reproduction, and lifespan. Moreover, our findings demonstrate that carbohydrate accelerates aging and impairs late-life physiology and behaviour. These results have implications for understanding the role of diet in shaping the life history traits of organisms. Notably, we detected distinct diet-induced variations in gene expression. Further in-depth investigations of the molecular pathways, such as Insulin signaling (IIS), target of rapamycin (TOR) signaling, and related pathways involved in energy homeostasis will provide us the key to understand underlying mechanisms behind the observed diet-induced plasticity in life-history, physiology, and behaviour. Such understanding holds the potential for devising therapeutic strategies targeting age-related diseases.

## Acknowledgements

We thank Prof. NG Prasad, IISER Mohali and his student Dr Rohit Kapila for providing us the outbred MB_1-4_ populations, which is the parental population of the experimental population used here. AM, CS and NK acknowledge the funding support from the Research and Development Office (RDO), Ashoka University. CM acknowledges the funding support from the Department of Biotechnology (DBT), India through the Junior Research Fellowship (JRF). We thank Prof. Sanjeev Galande for providing laboratory reagents and Prof. LS Shashidhara for access to lab equipment. ST acknowledges the support of DBT/Wellcome Trust India Alliance Early Career Fellowship (#IA/E/18/1/504347) for research funding and Ashoka University for the infrastructure.

## Supplementary Online Materials

### Text S1: Wing length measurement protocol

To understand the effect of diet on body size, we measured the wing size of the flies as a proxy. On the 12^th^ day post egg collection, flies were separated under CO_2_ anesthesia and arbitrarily one of the two wings were clipped off for 30 individuals per sex. The wings were then mounted on a microscopic slide using a drop of polyethylene glycol (PEG) and imaged. The length of the vein between distance between the anterior cross vein (ACV) and the third longitudinal vein (L3) was measured using ImageJ as a standard metric for wing size.

**Table S1.** Recipes to make isocaloric diets (714KCal). The flies get carbohydrates from sugar (energy = 3.87KCal/gm) and corn (energy = 3.33KCal/gm) whereas they get proteins and carbohydrates from yeast (energy = 3.54kCal/gm). The P:C ratio has been calculated considering the total protein and the total carbohydrate that has been contributed by each diet component.

| **P:C Ratio** | **0.2** | **0.25** | **0.4** | **0.55** | **0.7** |
| --- | --- | --- | --- | --- | --- |
| **Water (L)** | **1** | **1** | **1** | **1** | **1** |
| **Corn (g)** | **50** | **50** | **50** | **50** | **50** |
| **Sugar (g)** | **90.8** | **79.7** | **50** | **29** | **10.3** |
| **Yeast (g)** | **55.4** | **67.6** | **100** | **123** | **143.5** |
| **Agar (g)** | **15** | **15** | **15** | **15** | **15** |
| **Extra water (ml)** | **200** | **200** | **200** | **200** | **200** |
| **Benzoate (ml)** | **40** | **40** | **40** | **40** | **40** |
| **Propionic acid (ml)** | **20** | **20** | **20** | **20** | **20** |

\*Per gram, yeast contains 0.37g of carbohydrates and 0.48g of protein, while corn contains 0.78g of carbohydrates and 0.07g of protein.

The following tables are attached as separate Excel files for better data representations.

**Table S2. Complete List of Expressed Transcripts Across Samples**

**Table S3. Differential Transcript Expression Statistics: P:C = 0.25 Diet vs. Baseline (P:C = 0.4) Diet**

**Table S4. Differential Transcript Expression Statistics: P:C = 0.55 Diet vs. Baseline (P:C = 0.4) Diet**

**Table S5. Differential Transcript Expression Statistics: P:C = 0.7 Diet vs. Baseline (P:C = 0.4) Diet**

**Table S6. GO Analysis of Transcripts Upregulated in P:C = 0.25 Diet vs. Baseline (P:C = 0.4) Diet.**

**Table S7. GO Analysis of Transcripts Upregulated in P:C = 0.7 Diet vs. Baseline (P:C = 0.4) Diet.**

**Table S8.** Effect size of differential mean locomotor activity of old male flies reared on experimental diets.

| **Groups** | **Cohen's *d*** | **Inference** |
| --- | --- | --- |
| P:C 0.2 - P:C 0.25 | 0.2 | Small |
| P:C 0.2 - P:C 0.4 | 0.38 | Small |
| P:C 0.2 - P:C 0.55 | 0.15 | n.d. |
| **P:C 0.2 - P:C 0.7** | **0.57** | **Medium** |
| **P:C 0.25 - P:C 0.4** | **0.58** | **Medium** |
| P:C 0.25 - P:C 0.55 | 0.35 | Small |
| **P:C 0.25 - P:C 0.7** | **0.82** | **Large** |
| P:C 0.4 - P:C 0.55 | 0.24 | Small |
| P:C 0.4 - P:C 0.7 | 0.15 | n.d. |
| P:C 0.55 - P:C 0.7 | 0.41 | Small |

$d < 0.2$ : n.d. or no difference, $< d < 0.5$ : small, $0.5 < d < 0.8$ : medium), $d > 0.8$ : large

**Table S9.** Analysis of Deviance Table (Type III tests) comparing survival curves in lifespan assay using Cox-Proportional hazard model followed by Wald Test.

|  | Df | Chisq | Pr(>Chisq) |  |
| --- | --- | --- | --- | --- |
| Food | 4 | 68.5285 | 4.640e-14 | *** |
| Sex | 1 | 0.4473 | 0.5036 |  |
| Food:Sex | 4 | 36.3192 | 2.487e-07 | *** |

Signif. codes: 0 '\*\*\*' 0.001 '\*\*' 0.01 '\*' 0.05 '.' 0.1 ' ' 1

**Figure S1.**
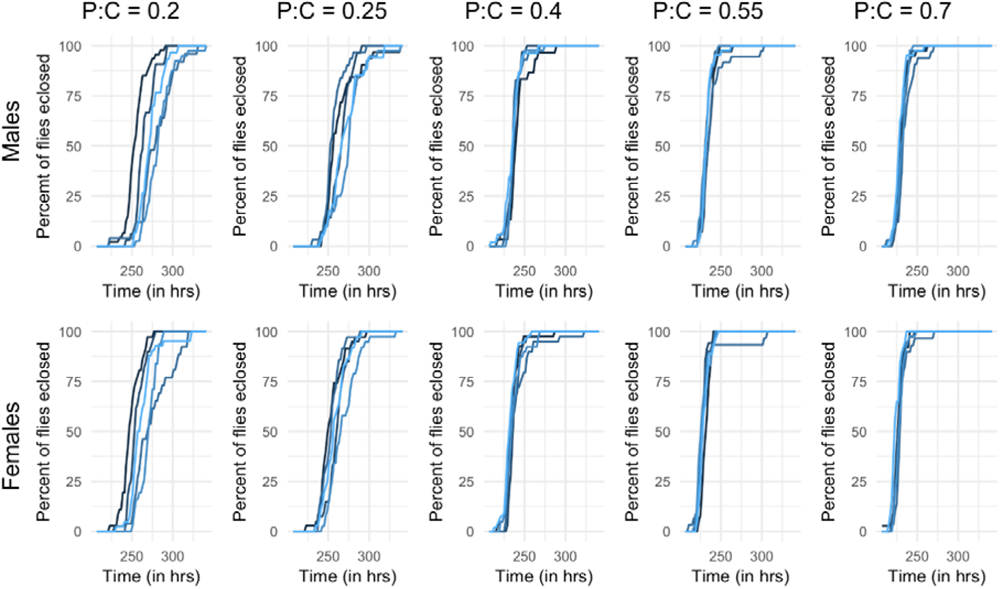
Mean percentage of flies of A) female and B) male flies that eclosed over time which were reared on different diets. Each line represents a replicate of the experiment.

**Figure S2.**
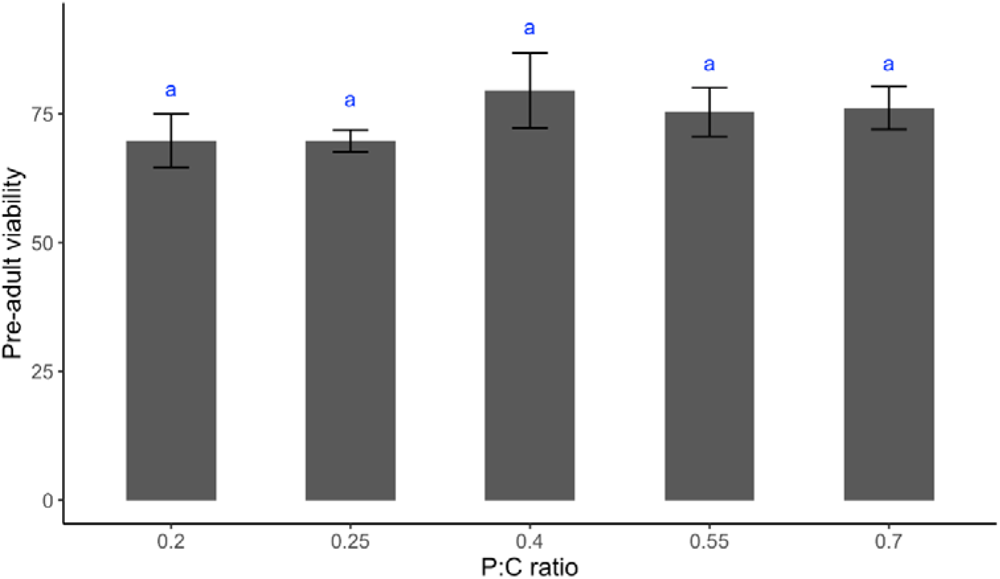
Mean (±SD) pre-adult viability of flies being reared on different diets. The bar plot represents the pre-adult viability of the flies being reared on diets with different P:C ratios. The error bar represents the standard deviation of the pre-adult viability data.

**Figure S3.**
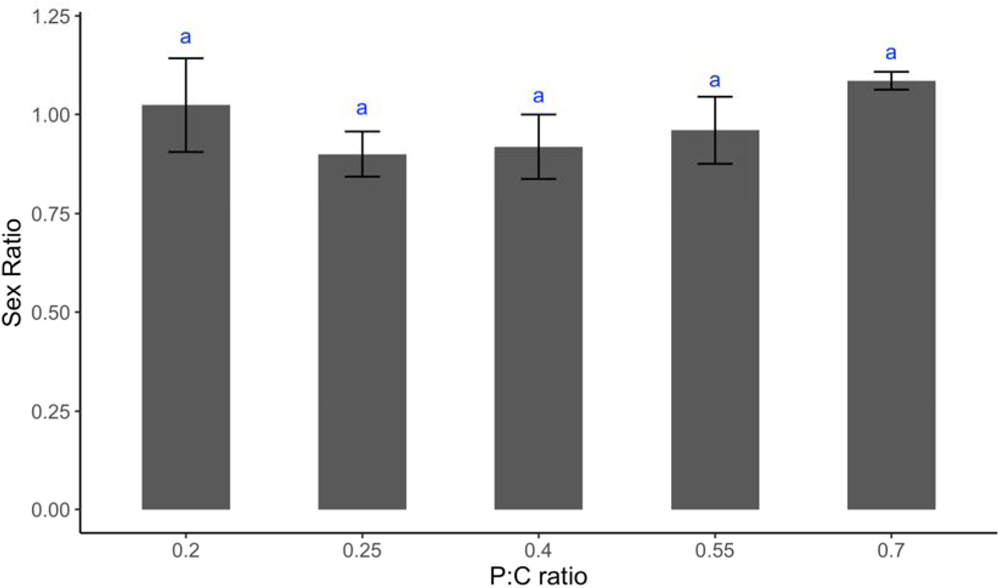
Mean (± SD) sex ratio of flies being reared on different diets. The bar plot represents the sex ratio of the flies being reared on diets with different P:C ratios. The error bar represents the standard deviation of the sex ratio data.

**Figure S4.**
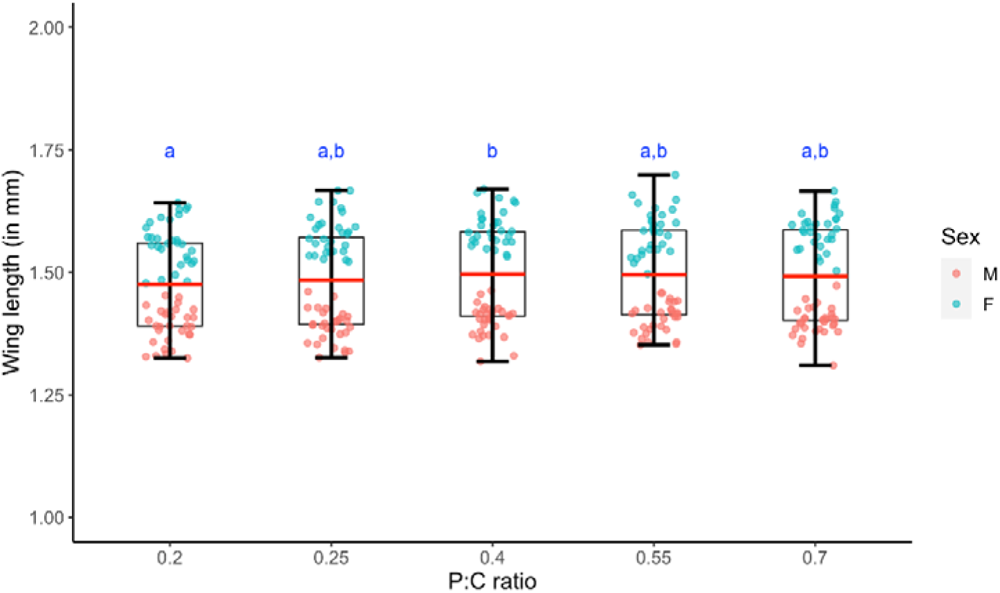
Wing length as a proxy for body size of flies that were reared on different diets. The box plot represents the wing length of the male and female flies, which were reared on diets with different P:C ratios. The bold red line represents the mean wing length of combined male and female data. The cyan and red scatter points represent wing length of the individual female and male flies, respectively.

**Figure S5.**
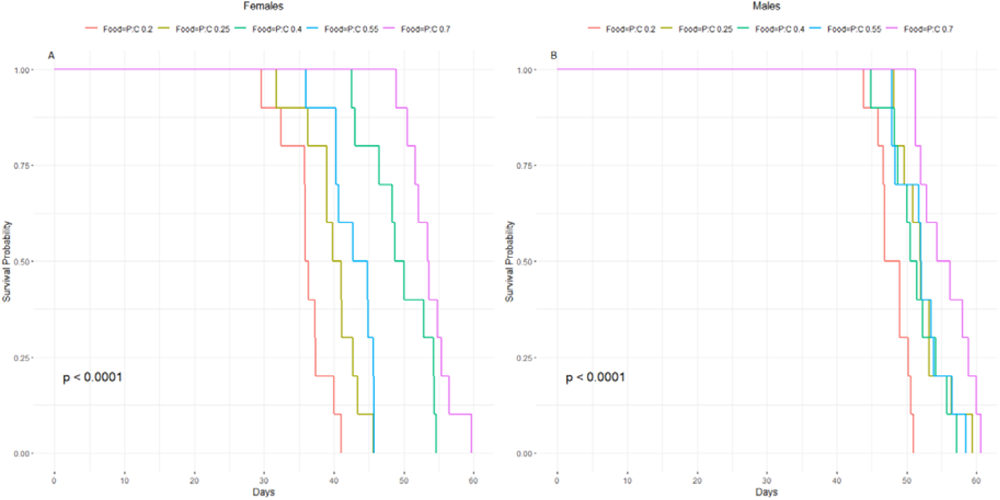
Mean survival probability of (A) female and (B) male flies over days that are reared on different diets. Each line represents the diet on which the flies were reared on.

**Figure S6.**
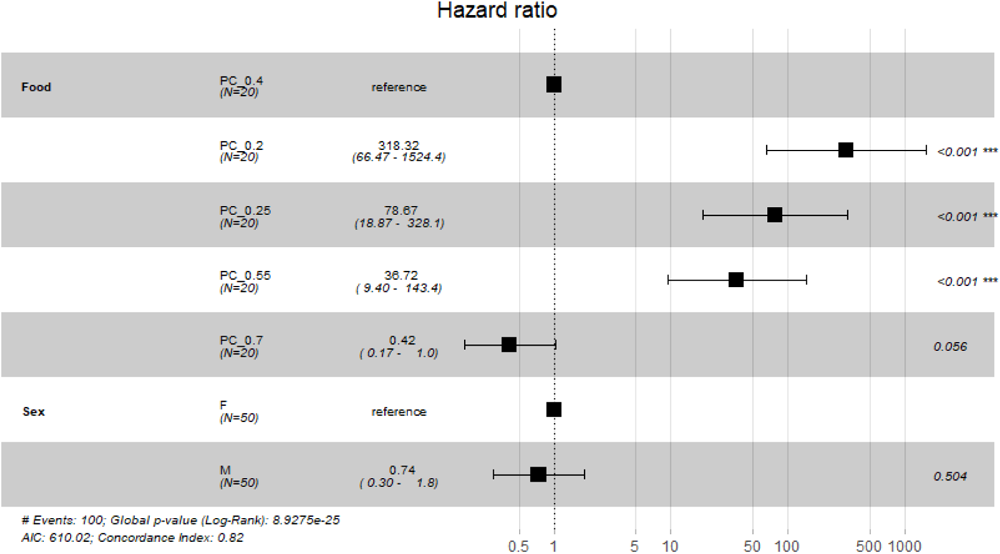
Hazard ratios for the impact of diet composition and sex on lifespan. This forest plot illustrates the hazard ratios for survival as influenced by different protein-to-carbohydrate (P:C) ratios and sex. The reference category for diet is P:C ratio of 0.4, and for sex, it is female (F). Each square represents the hazard ratio point estimate for a given category, with horizontal lines denoting 95% confidence intervals. The dashed vertical line denotes a hazard ratio of 1, indicating no effect. The numbers on the right-hand side indicates statistical significance (p-value). Statistically significant differences (p < 0.05) are marked with asterisks. The global p-value from the Log-Rank test and model fit statistics including Akaike Information Criterion (AIC) and Concordance Index are provided at the bottom of the plot.

**Figure S6:**
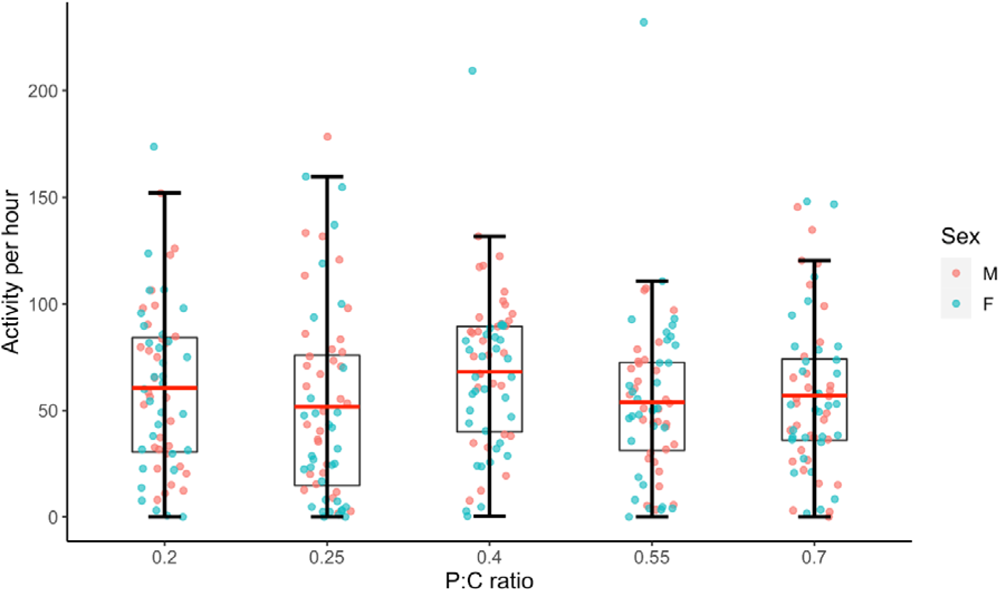
Mean (± SD) activity of flies per hour that are reared on different diets. The box plot represents the activity per hour of the male and female flies, which were reared on diets with different P:C ratios. The bold red line represents the mean activity of combined male and female data. The cyan and red scatter points represent activity of the individual female and male flies, respectively.

**Figure S8.**
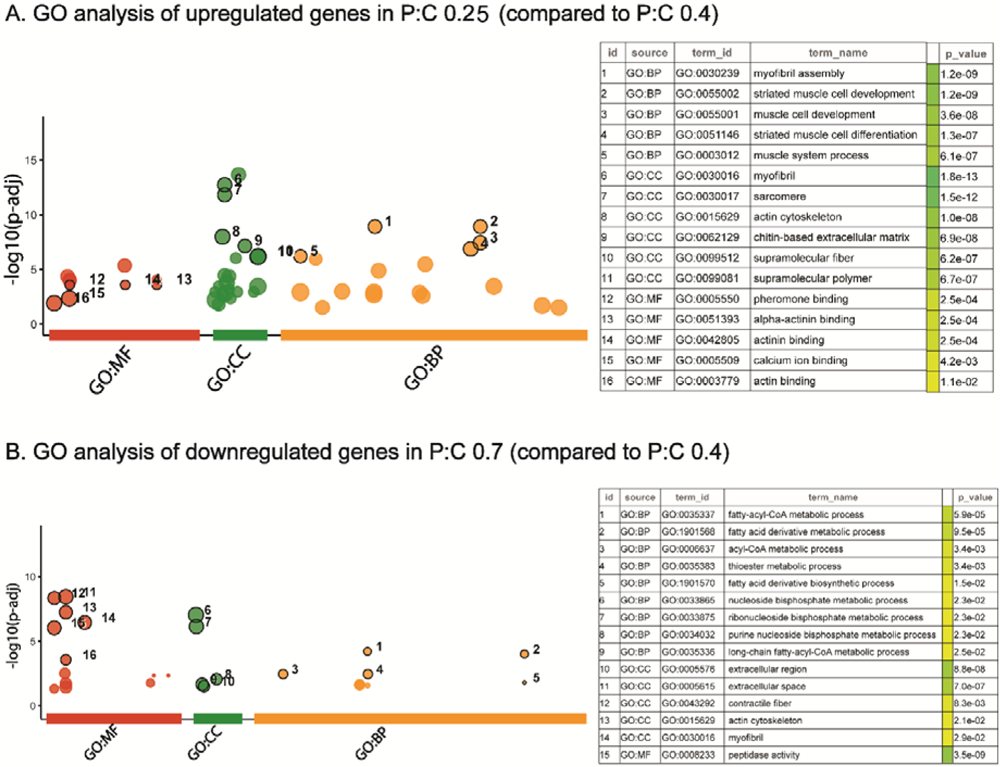
Gene Ontology (GO) analysis of gene expression in response to dietary protein-to-carbohydrate ratios. (A) Upregulated gene GO terms in P:C 0.25 vs. P:C 0.4, and (B) downregulated gene GO terms in P:C 0.7 vs. P:C 0.4 are presented here. Bubble size indicates term size. The y-axis reflects the significance (-log10(p-adj)) across biological processes (BP), cellular components (CC), and molecular functions (MF). Key enriched terms are labeled numerically and detailed in the adjacent table, which lists their GO IDs, terms, and p-values.

